# Macrophages control pathological interferon responses during viral respiratory infection

**DOI:** 10.1101/2023.12.16.572019

**Authors:** Daisy A. Hoagland, Patricia Rodríguez-Morales, Alexander O. Mann, Shuang Yu, Alicia Lai, Alan Baez Vazquez, Scott D. Pope, Jaechul Lim, Shun Li, Xian Zhang, Ming O. Li, Ruslan Medzhitov, Ruth A. Franklin

## Abstract

Antiviral immune mediators, including interferons and their downstream effectors, are critical for host defense yet can become detrimental when uncontrolled. Here, we identify a macrophage-mediated anti-inflammatory mechanism that limits type I interferon (IFN-I) responses. Specifically, we found that cellular stress and pathogen recognition induce Oncostatin M (OSM) production by macrophages. OSM-deficient mice succumbed to challenge with influenza or a viral mimic due to heightened IFN-I activation. Macrophage-derived OSM restricted excessive IFN-I production by lung epithelial cells following viral stimulation. Furthermore, reconstitution of OSM in the respiratory tract was sufficient to protect mice lacking macrophage-derived OSM against morbidity, indicating the importance of local OSM production. This work reveals a host strategy to dampen inflammation in the lung through the negative regulation of IFN-I by macrophages.

**One-Sentence Summary:** Type I interferons induced by viral stimuli are negatively regulated by macrophage-derived Oncostatin M.

## Main Text

Inflammation requires precise regulation to limit tissue damage and preserve organ function. This is exemplified in viral respiratory infections which require rapid and robust immune responses for viral clearance while maintaining efficient lung function (*1*). Upon exposure to respiratory viruses, the host mounts a complex and coordinated cascade of responses to restrict and eliminate infection (*2, 3*). Viral encounter is first sensed by pattern recognition receptors that detect conserved pathogen-associated molecular patterns (PAMPs) at the plasma membrane, in the cytosol, or in endosomal compartments (*4*). These recognition pathways activate production of numerous chemokines and cytokines to further activate innate and adaptive immunity, including members of the evolutionarily conserved family of interferons (*5*). Type I interferons (IFN-I) are essential antiviral cytokines produced by both immune and nonimmune cell types (*6*). While critical for protection against viral infection, inflammatory responses driven by IFN-I can become pathological, causing cell death, tissue damage, and impaired repair (*7*–*11*). Therefore, IFN-I production must be tightly controlled to allow effective antiviral defense while preventing immunopathology.

Macrophages are innate immune cells with key roles in activating inflammatory responses to defend the host against pathogens. Macrophages also orchestrate the resolution of inflammation and maintenance of tissue homeostasis (*12*–*16*). Our knowledge of negative regulators of inflammation is currently limited to a few mediators, including IL-10, TGF-?, and glucocorticoids. Here we uncover a new mechanism by which macrophages suppress IFN-I production through the cytokine Oncostatin M (OSM) following recognition of viral stimuli. In the absence of macrophage-derived OSM, lung epithelial cell production of IFN-I is exacerbated, and mice succumb to influenza infection, or treatment with the viral mimic, polyinosinic-polycytidylic acid [poly(I:C)]. These findings suggest that macrophages limit the negative consequences of robust host defense responses following respiratory viral infection to promote survival.

### OSM is induced following influenza infection and is necessary for survival

Macrophages are key players in the regulation of inflammation and restoration of tissue homeostasis following infection. We hypothesized that in response to stress and damage, macrophages produce dedicated signals to restore tissue homeostasis (Fig. 1A). We identified the cytokine OSM as induced in macrophages by a variety of stressors including heat shock and ER stress, and PAMPs including LPS and poly(I:C) (Fig. 1B and fig. S1A). OSM, a cytokine produced predominantly by myeloid cells and T cells, signals through glycoprotein 130 (gp130) and OSM receptor (OSMR) in mice. Studies have suggested that OSM can exert both pro- and anti-inflammatory effects in a context-dependent manner (*17, 18*), but the biological mechanisms underlying these differences remain obscure. Given the rapid synthesis of OSM in response to stress and PAMPs *in vitro*, we utilized a murine model of Influenza A virus (IAV) infection, which induces cellular stress and generates viral-derived PAMPs (*19*). Upon infection of animals with a sublethal dose of IAV (A/WSN/1933 (H1N1); fig. S2A), we observed local OSM production in the airways, as assessed by RNA expression in whole lung and protein expression in bronchoalveolar lavage fluid (BALF), but not in blood plasma (Fig. 1, C and D).

**Fig. 1.**
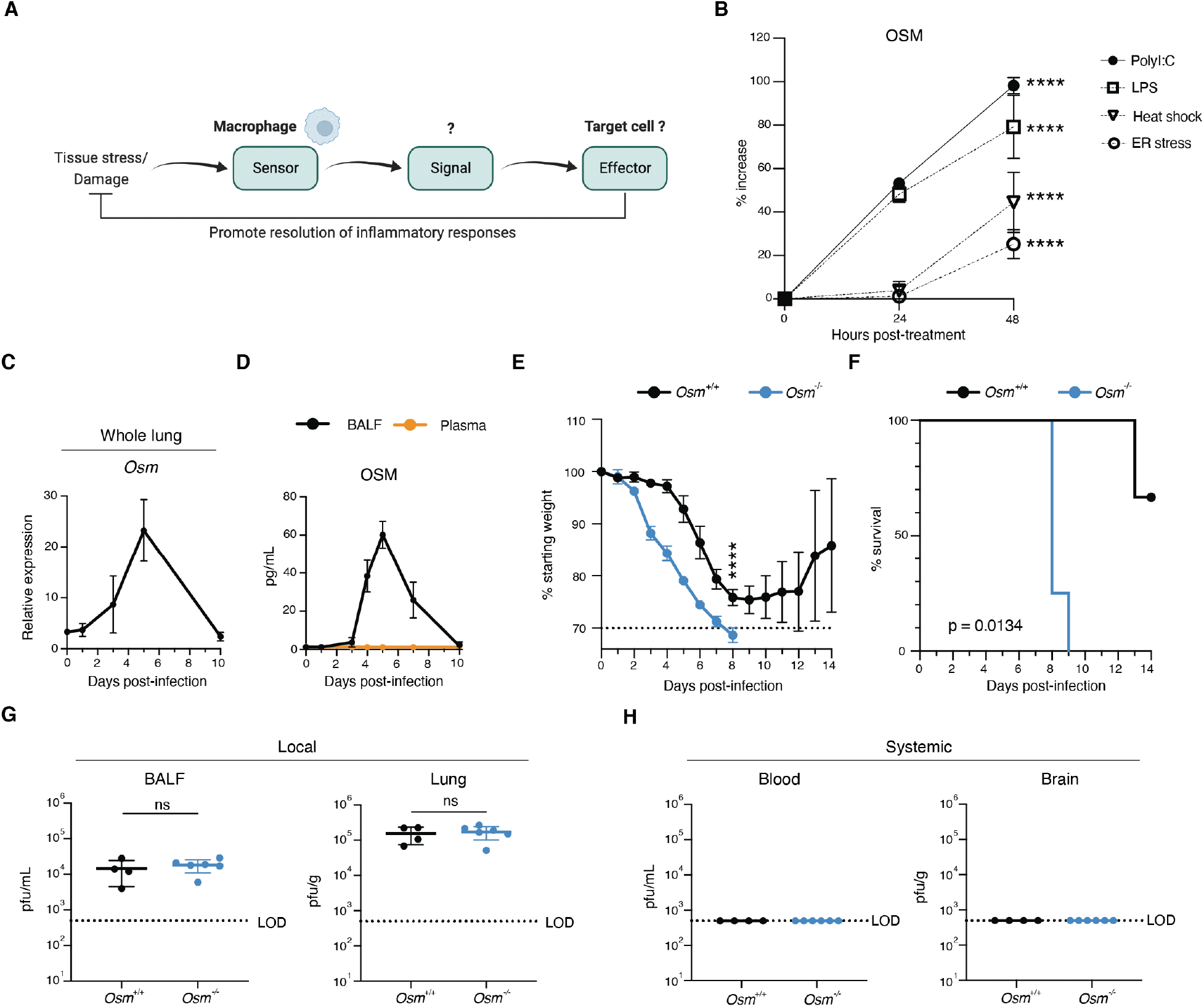
OSM is induced by viral infection and is required for host survival. (**A**) Illustration of overall hypothesis. (**B**) Bone-marrow derived-macrophages (BMDMs) were treated with indicated PAMPs or stressors, supernatant was collected at indicated time points, and protein levels were assessed by ELISA (n = 3). **(C** and **D)** Mice were infected intranasally with 225 PFU of A/WSN/1933(H1N1), and RNA from whole lung (C) and protein from BALF and plasma (D) were harvested at indicated time points (n = 3). (**E** and **F**) Percent initial body weight (E) and survival curve after infection (F) with 225 PFU of A/WSN/1933(H1N1), (n = 3 - 4). (**G** and **H**) Plaque forming units (PFU) detectable in BALF, lung, blood, and brain at 6 days post infection (dpi) (n = 4 – 6). Relative expression calculated as (2^-ΔCt^)*1000. All data are representative of at least two independent experiments. Error bars indicate standard deviation (SD), ****p ≤ 0.0001, ns = not significant, (unpaired Student’s t test for (B) and (G), two-way ANOVA up to day 8 for (E), log-rank Mantel-Cox test for (F). LOD = level of detection.

The induction of OSM during IAV infection led us to hypothesize that it may play an important role in the immune response to viral infection. To address this question, we established an *Osm*-deficient mouse line and infected *Osm*^*+/+*^ and *Osm*^*-/-*^ mice with IAV. Strikingly, *Osm*^-/-^ mice displayed increased susceptibility to IAV infection, as observed by increased morbidity and mortality (Fig. 1, E and F). We next investigated whether increased pathogen burden might be responsible for the survival defect observed in *Osm*^-/-^ mice. Upon examining viral load in the lung and BALF six days post-infection, we observed that *Osm*^-/-^ mice exhibited no differences in viral titer or viral transcripts in the respiratory tract when compared to littermate *Osm*^*+/+*^ mice (Fig. 1G and fig. S2B). In addition, we evaluated viral load in the brain and blood to investigate whether viral dissemination was responsible for the increased mortality in *Osm*^-/-^ mice and found no detectable virus in these tissues (Fig. 1H and fig. S2B). Together, these data suggest that antiviral defense mechanisms are intact in *Osm*^-/-^ mice.

### *Osm*-deficient mice display increased levels of IFN-I during acute infection

To investigate differences in gene expression that could account for the survival phenotype following IAV infection, we performed bulk RNA-sequencing on whole lung RNA from *Osm*^-/-^ and *Osm*^*+/+*^ mice at baseline, and three or seven days post-IAV infection (Fig. 2A). We then carried out pathway enrichment analysis using Enrichr (*20*) on genes that were upregulated in IAV-infected *Osm*^-/-^ mice as compared to IAV-infected *Osm*^*+/+*^ mice (fig. S3A). Three days after infection, the highest enrichment score from this gene list was for interferon alpha/beta signaling (Fig. 2B). Furthermore, Molecular Signatures Database (MSigDB) (*21*) scoring of the IFNα signature expression across individual samples revealed enhanced IFNα response signature genes in *Osm*^-/-^ mice (Fig. 2C and fig. S3B). The majority of the genes in this signature represent interferon-stimulated genes (ISGs), the effector molecules downstream of interferons (*22*). While upregulation of an IFN-I signature was anticipated in infected mice compared to uninfected controls, upregulation of IFN-I signaling was also observed in infected *Osm*^-/-^ mice compared to infected *Osm*^*+/+*^mice, indicating more robust IFN-I signaling in the absence of OSM following IAV infection.

**Fig. 2.**
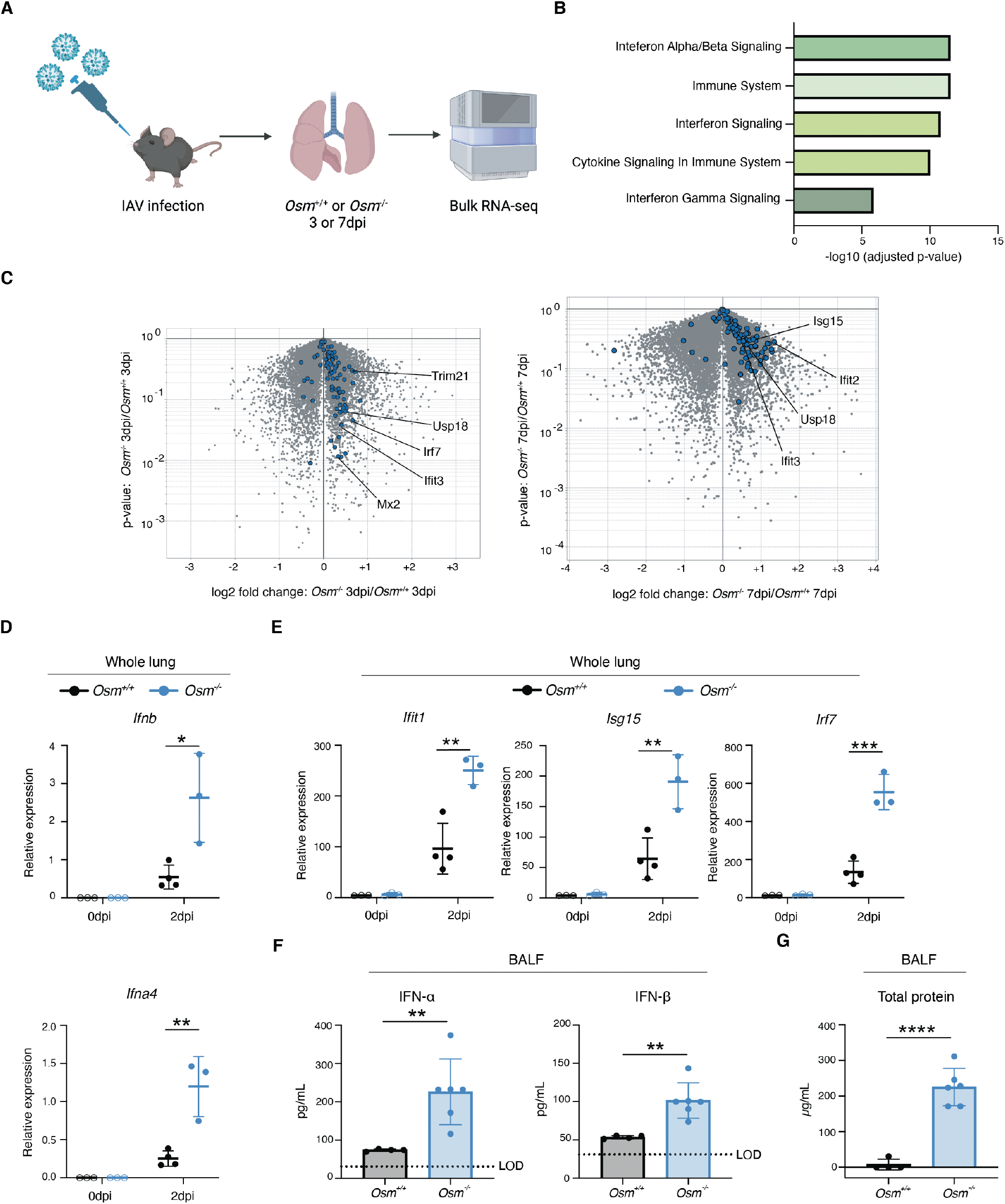
OSM deficiency leads to elevated levels of type I IFN during IAV infection. (**A**) Schematic of experimental design for (B) and (C). Mice were infected with 225pfu of A/WSN/1933(H1N1) and whole lung RNA was harvested three or seven days post-infection for bulk RNA-sequencing. (**B** and **C**) Transcriptional analysis of whole-lung gene expression (n = 2). Pathway enrichment analysis performed using Enrichr on genes that were at least 0.5 log2 fold change upregulated in both *Osm*^*+/+*^ and *Osm*^*-/-*^ mice at 3dpi versus uninfected, and 0.25 log2 fold change more differentially upregulated in IAV-infected *Osm*^-/-^ mice as compared to *Osm*^*+/+*^ mice. Gene expression pathways were retrieved from the Reactome Database (B). Volcano plot illustrating expression of MSigDB hallmark interferon alpha response signature genes between *Osm*^*+/+*^ and *Osm*^*-/-*^ mice at 3dpi and 7dpi (C). (**D** and **E**) RNA from whole lung was harvested at indicated time points. (**F**) Protein from BALF harvested at 2dpi (n = 3 - 4; representative of two independent experiments). (**G**) Total protein in BALF at 2dpi (n = 4 – 6). IFN-α ELISA for IFN-α2 subtype. Relative expression calculated as (2^-ΔCt^)*1000. Error bars indicate SD, *p ≤ 0.05, **p ≤ 0.01, ***p ≤ 0.001, ****p ≤ 0.0001, (unpaired Student’s t test for (D), (E), and (G); Mann-Whitney U test for (F)).

To determine if the increase in ISG expression in infected mice lacking *Osm* correlated with increased transcription of IFN-I, we harvested BALF and whole lung RNA from *Osm*^-/-^ and *Osm*^*+/+*^ mice at baseline and two days after infection. Analysis of whole lung RNA by RT-qPCR revealed increased levels of IFN-I transcripts, including *Ifnb* and *Ifna4* (Fig. 2D). This was accompanied by an increase in ISG expression, exemplified by elevated *Ifit1, Isg15*, and *Irf7* transcripts (Fig. 2E). IAV-infected *Osm*^-/-^ mice also showed increased levels of IFN-I in BALF compared to *Osm*^*+/+*^ mice (Fig. 2F). Notably, there were no differences in IFN-I transcripts, ISG transcripts, or IFN-I protein levels at baseline between *Osm*^-/-^ and *Osm*^*+/+*^ mice, indicating these differences in IFN-I must be driven by the presence of viral stimuli (Fig. 2, D and E and fig. S3C). IAV-infected *Osm*^-/-^ mice also displayed increased levels of total protein in BALF as compared to infected *Osm*^+/+^ mice, indicating a loss of lung barrier integrity (Fig. 2G), a known hallmark of interferon-mediated immunopathology (*9*). Based on these observations, we hypothesized that enhanced IFN-I production in *Osm*^*-/-*^ mice underlies their increased morbidity and mortality.

### Macrophage-derived OSM is required to survive challenge with a viral mimic

To further investigate the potential regulation of IFN-I by OSM, we utilized an additional *in vivo* model consisting of intratracheal (i.t.) delivery of a potent type I interferon-inducing stimulus, poly(I:C) (Fig. 3A). Poly(I:C) mimics a double-stranded RNA viral replication product and stimulates TLR3 and RIG-I/MDA5 pathways (*23*). Leveraging this model allowed us to eliminate confounding factors introduced by live virus, such as viral replication, while still mounting innate immune responses and generating epithelial damage (*24*). Similar to IAV infection, we observed that OSM concentrations increased locally in BALF in response to i.t. poly(I:C) treatment, but not systemically in blood plasma (Fig. 3, B and C and fig. S4A). We also observed *Osm* induction in lung monocytes/macrophages upon treatment with poly(I:C), indicating their relevance as a source of OSM *in vivo* in this model (fig. S4, B and C). Furthermore, *Osm*^*-/-*^ mice were highly susceptible to poly(I:C) treatment (Fig. 3, D and E) as opposed to *Osm*^+/+^ mice, recapitulating the survival and morbidity phenotypes observed during IAV infection. The observation that mice did not survive exposure to viral PAMP emphasizes the inability of OSM-deficient mice to prevent severe outcomes from what is typically transient activation of innate immunity. Moreover, we observed this survival phenotype in both male and female mice, suggesting a conserved role for OSM across sexes (fig. S5, A and B).

**Fig. 3.**
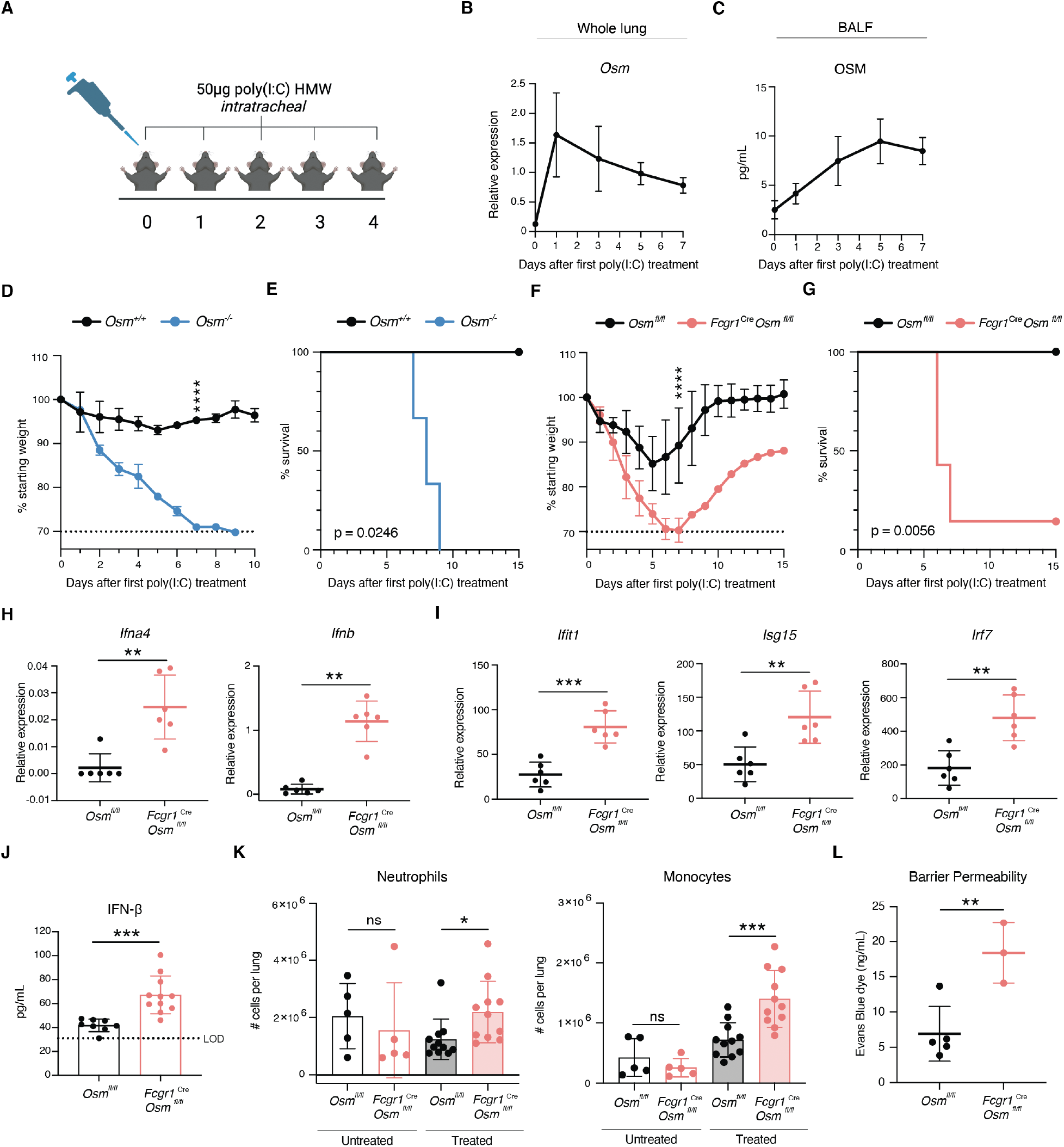
Macrophage-derived OSM is required for survival during poly(I:C) challenge. (**A**) Schematic of experimental design. Mice were treated intratracheally (i.t.) with poly(I:C) (50μg) daily for five consecutive days and monitored daily. (**B**) and (**C**) Whole lung RNA and BALF were collected at indicated time points (n = 4 - 5). (**D – G**) Mice were treated i.t with poly(I:C) and monitored for body weight (D) and (F) and survival (E) and (G) (n = 3 - 7). (**H** – **J**) Whole lung RNA (H) and (I) and BALF (J) were collected at day 7 after the first poly(I:C) treatment (n = 3 - 5). (**K**) Lung neutrophils and monocytes were quantified by flow cytometry at baseline and at day 7 after the first poly(I:C) treatment (n = 5 - 11; pooled from three independent experiments for poly(I:C) treated mice). (**L**) Lung barrier permeability Evans blue dye (EBD) assay of BALF at day 7 after the first poly(I:C) treatment (n = 3 – 5). A combination of female and male mice was used for experiments in this figure. Relative expression calculated as (2^-^ ΔCt)*1000. (B), (C), (D), (E), (F), (G), (H), (I), (J) and (K) are representative of at least two independent experiments. Error bars indicate SD, *p ≤ 0.05, **p ≤ 0.01, ***p ≤ 0.001, ****p ≤ 0.0001, ns = not significant (two-way ANOVA up to day 7 for (D) and (F), log-rank Mantel-Cox test for (E) and (G), unpaired Student’s t test or Mann-Whitney U test for (H), (I), (J), (K), and (L).

To specifically interrogate the importance of macrophage-derived OSM upon poly(I:C) challenge, we developed a macrophage-specific conditional knockout mouse model by crossing *Osm*^*fl/fl*^ and *Fcgr1*^Cre^ mouse lines (*25*). We validated the loss of *Osm* in sorted macrophages at baseline (fig. S6, A and B) and analysis of *Osm* expression in whole lung following poly(I:C) challenge revealed that macrophages accounted for the majority of *Osm* expression during poly(I:C) challenge (fig. S6C). We observed that macrophage-specific deletion of *Osm* resulted in the same survival and morbidity phenotypes as whole-body *Osm*^-/-^ mice in response to poly(I:C) (fig. S3, F and G). These data reveal that macrophage-derived OSM is required for survival during viral PAMP-mediated lung inflammation.

To dissect the impact of macrophage-derived OSM on IFN-I regulation during the inflammatory response to viral stimuli, we investigated the differences in IFN-I and ISG induction in *Fcgr1*^Cre^*Osm*^*fl/fl*^ and *Osm*^*fl/fl*^ mice three days after completion of our poly(I:C) treatment regimen. *Fcgr1*^Cre^*Osm*^*fl/fl*^ mice displayed increased expression of IFN-I and ISGs in the lung (Fig. 3, H, I and J), further supporting the hypothesis that OSM negatively regulates IFN-I responses. This increase in IFN-I in the *Fcgr1*^Cre^*Osm*^*fl/fl*^ mice was accompanied by enhanced inflammation, as evidenced by monocyte and neutrophil infiltration (Fig. 3K and fig. S7). Previous studies have demonstrated interferon-driven disruption of lung epithelial barrier function (*9, 24*). Therefore, we tested whether mice lacking macrophage-derived OSM would exhibit a loss of epithelial cells and barrier integrity. Indeed, *Fcgr1*^Cre^*Osm*^*fl/fl*^ mice showed a reduction in lung epithelial cells, both in total number and fraction of non-immune cells (fig. S8, A and B). Importantly, this decrease was also evident for type II pneumocytes (ATIIs) (fig. S8, A and C), a key epithelial progenitor population, suggesting alveolar damage (*26*). To further investigate disruption of barrier integrity in the lung, we intraperitoneally injected Evans blue dye (EBD) into mice following poly(I:C) treatment and assessed the EBD levels in BALF. EBD levels were increased in *Fcgr1*^Cre^*Osm*^*fl/fl*^ mice, accompanied by increased total protein levels (Fig. 3L and fig. S8D). Together, these data indicate that macrophage-derived OSM dampens innate immune responses and promotes barrier integrity in response to viral stimuli to promote survival.

### Blockade of IFN-I signaling or administration of recombinant OSM rescues *Osm*-deficient mice from poly(I:C)-driven morbidity

To directly test the hypothesis that OSM protects mice from succumbing to poly(I:C) treatment through the negative regulation of IFN-I responses, we administered intravenous (i.v.) IFN-α/β receptor antibodies (anti-IFNAR1) to *Fcgr1*^Cre^ *Osm*^*fl/fl*^ and *Osm*^*fl/fl*^ mice during treatment with poly(I:C). Upon IFNAR1 neutralization, *Fcgr1*^Cre^ *Osm*^*fl/fl*^ mice exhibited no differences in morbidity or mortality compared to anti-IFNAR1 treated WT mice (Fig. 4A and fig. S9A). This demonstrates that IFN-I signaling is required for the mortality observed in mice lacking macrophage-derived OSM following treatment with poly(I:C). Furthermore, we administered murine recombinant OSM (rOSM) intratracheally to *Fcgr1*^Cre^ *Osm*^*fl/fl*^ and *Osm*^*fl/fl*^ mice in combination with our poly(I:C) treatment regimen. Reconstitution of OSM in the respiratory tract lessened morbidity in *Fcgr1*^Cre^ *Osm*^*fl/fl*^ mice (Fig. 4B and fig. S9B). This indicates that local OSM signaling in the respiratory tract is sufficient to ameliorate pathological effects resulting from OSM deficiency in macrophages.

**Fig. 4.**
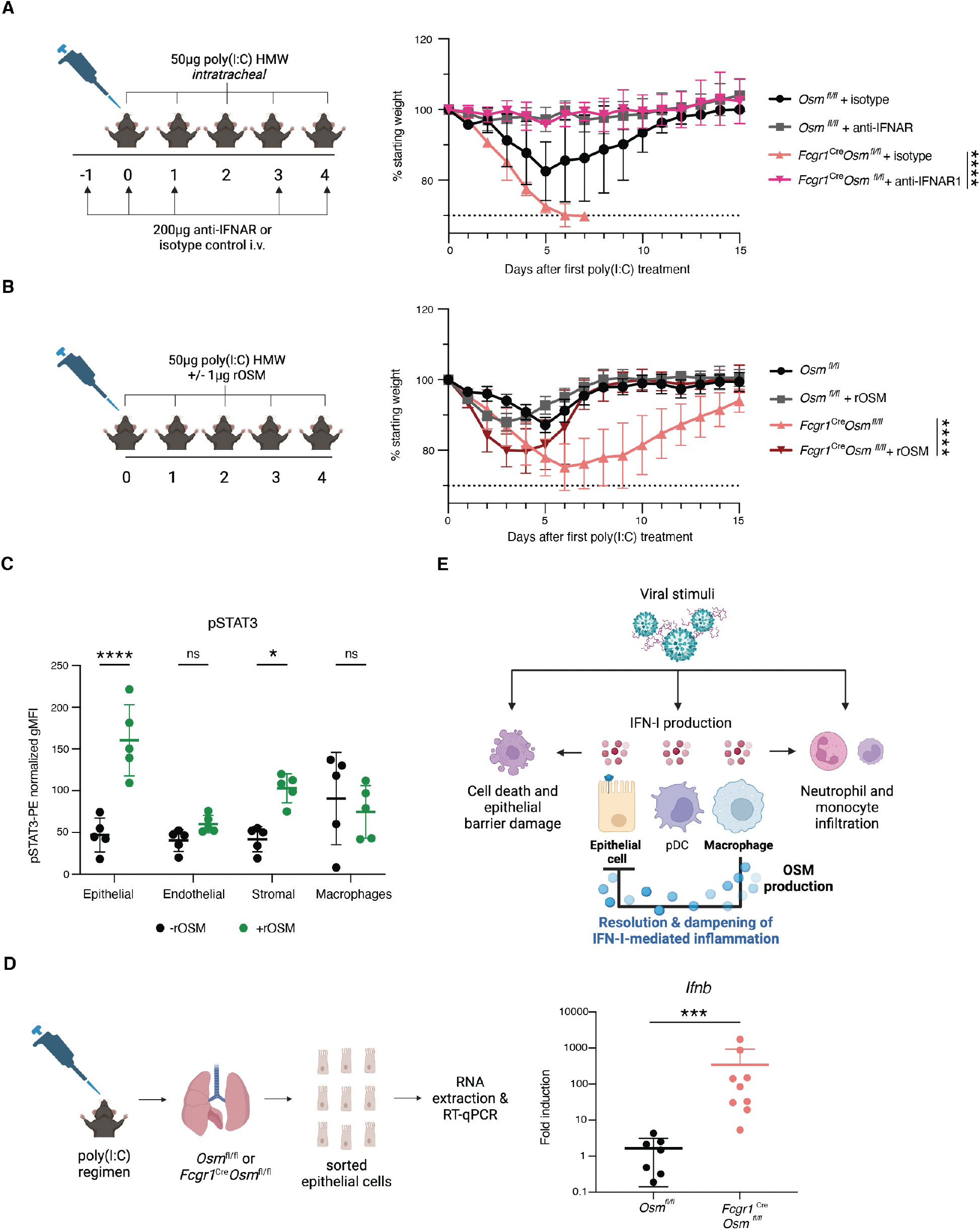
Type I IFN signaling blockade restores survival in poly(I:C)-treated mice. (**A**) Mice were treated intravenously (i.v.) with either 200μg of anti-mouse IFNAR-1 or isotype control antibody diluted in 100μl PBS on day -1, day 0, day 1, day 3, and day 4 after the first poly(I:C) treatment. Mice were weighed daily (n = 3 - 4). (**B**) Mice were treated i.t. with 1µg of murine rOSM in addition to 50µg of poly(I:C) daily for five consecutive days, and weighed daily (n = 4 - 5). (**C**) Cells isolated from digested lungs from WT mice were stimulated *ex vivo* with rOSM (100ng/mL) for 30 minutes and stained for surface markers and intracellular pSTAT3. Summary representation of pSTAT3 geometric mean fluorescence intensity (gMFI) in indicated populations following stimulation. pSTAT3-PE fluorescence signal was normalized per population by subtracting the PE signal in pSTAT3 fluorescence minus one controls (FMO) from the stained sample gMFI values. Epithelial cells were defined as EpCAM^+^ CD45^-^ CD31^-^, endothelial cells as CD31^+^ EpCAM^-^ CD45^-^, stromal as CD45^-^ CD31^-^ CD45^-^, and macrophages as CD45^+^ CD64^+^, all gated on viable singlets (n = 5; representative of two independent experiments). (**D**) RT-qPCR for *Ifnb* in fluorescence activated cell sorting (FACS)-sorted epithelial cells at day 7 after the first poly (I:C) treatment (n = 7 - 8). Data pooled from two individual experiments. (**E**) Model of OSM-mediated inhibition of IFN-I production in epithelial cells. (A) and (B) are representative of two independent experiments. Relative expression calculated as (2^-ΔCt^)*1000. Error bars indicate SD, *p ≤ 0.05, ***p ≤ 0.001, ****p ≤ 0.0001, ns = not significant, (unpaired Student’s t test for (C), Mann-Whitney U test for (D), Two-way ANOVA up to day 6 for (A) and up to day 15 for (B).

### Macrophage-derived OSM negatively regulates IFN-I production by epithelial cells

To elucidate which cell populations in the lung OSM may be acting on to suppress IFN-I, we examined *Osmr* expression in various cell types in poly(I:C) treated and untreated *Fcgr1*^Cre^ *Osm*^*fl/fl*^ and *Osm*^*fl/fl*^ mice and did not observe differences in receptor expression across genotypes (fig. S10, A, B, and C). Receptor expression was highest in non-immune cells including epithelial cells, endothelial cells, and fibroblasts (fig. S10, A, B, and C). To further interrogate responsiveness to OSM, we digested whole lung and stimulated cells with rOSM *ex vivo*. Epithelial cells exhibited the most robust downstream response to OSM, as measured by STAT3 phosphorylation (pSTAT3) (*27*), with stromal cells responding to a lesser extent (Fig. 4C and fig. S10D). In contrast, there was not a significant change in pSTAT3 signal in endothelial cells or monocytes/macrophages (Fig. 4C). This suggested that epithelial cells may play a role in the direct sensing of OSM, leading to dampened IFN-I responses. To further test the responsiveness of airway epithelial cells to OSM, we transfected a human alveolar basal epithelial cell line (A549) with poly(I:C) and stimulated with rOSM. Human rOSM-treated A549 cells displayed decreased levels of ISGs including *IFIT1* and *IRF7* (fig. S10E), suggesting a direct role for OSM on IFN-I signaling in the epithelium.

To investigate OSM signaling in epithelial cells *in vivo*, we sorted epithelial cells from the lungs of *Fcgr1*^Cre^ *Osm*^*fl/fl*^ and *Osm*^*fl/fl*^ mice following poly(I:C) treatment (fig. S10A). We found that epithelial cells from mice lacking macrophage-derived OSM displayed increased transcription of *Ifnb* (Fig. 4D). This difference in *Ifnb* taken together with *Osmr* expression and pSTAT3 activation in epithelial cells reveals a previously undescribed communication circuit to constrain inflammatory responses. IFN-I is produced by a variety of cell types including epithelial cells, pDCs, and macrophages upon viral encounter in the lung (Fig. 4E). Robust IFN-I responses promote further inflammation, characterized by neutrophil and monocyte recruitment and collateral damage to the epithelial barrier (Fig. 4E). We propose a model in which macrophages not only participate in production of antiviral effectors but additionally produce OSM downstream of viral stimuli, limiting IFN-I transcript levels in epithelial cells, and thus preventing pathological IFN-I responses and promoting host survival (Fig. 4E).

## Discussion

Our work found that macrophages produce the cytokine OSM in response to viral stimuli to control pathological IFN-I responses. In the absence of OSM, we observed enhanced IFN-I activity in the lung, including dysregulated IFN-I and ISG induction in lung epithelial cells which resulted in a shift to pathological IFN-I responses and increased mortality. IFN-I responses have widespread effects as IFNAR is expressed ubiquitously across all cell types (*28*). In addition to protective responses, IFN-I can cause pathology due to proinflammatory activity and antiproliferative and proapoptotic functions (*7*–*10*). Studies have also demonstrated that both type I and type III interferons can play protective or pathological roles depending on the timing of their expression and the specific anatomical site within the respiratory tract where they are active (*29*). As such, macrophage-derived OSM dampens IFN-I responses in epithelial cells to promote a tissue state compatible with repair. This exemplifies the importance of epithelial-immune crosstalk, analogous to repair circuits described in other tissues (*30*–*32*).

Macrophages are both potent inducers of inflammation in response to infection and important in tissue repair and restoration of homeostasis (*15*). Many of these immunomodulatory functions occur in later stages of infection (*33, 34*). Interestingly, we found that OSM is induced in macrophages rapidly after detection of viral stimuli, suggesting that OSM is an early immune modulator. This swift OSM induction may ensure that IFN-I responses remain within a physiological range throughout the disease course to prevent pathology, and/or to regulate IFN-I in a particular region of the airway as viral infection progresses. Indeed, we found that i.t. rOSM administration rescues *Fcgr1*^Cre^ *Osm*^*fl/fl*^ mice from IFN-I-induced morbidity, suggesting that OSM is acting locally following viral stimuli.

Previous studies have described both pro- and anti-inflammatory roles for OSM (*35*–*37*). As *Osm* is induced in response to a variety of cues, it is possible that the tissue context of induction may shape the role of OSM. It is also possible that the negative regulation of IFN-I by OSM described in our study may explain some of these paradoxical findings. For example, in the context of intestinal inflammation, high levels of OSM were found in patients that were refractory to anti-TNF therapy (*36*). Similar findings were observed in mice, where OSM-deficiency resulted in decreased immune infiltration, epithelial and goblet cell disruption, and overall pathology in a model of anti-TNF resistant colitis (*36*). Perhaps these findings are due to indirect effects of OSM on IL-17-producing cells. IFN-I is known to constrain Th17 differentiation and IL-17A production (*38*), thus decreased colitis in the absence of OSM could be due to enhanced levels of IFN-I and decreased levels of IL-17. Indeed, *Il17a* transcripts remain high in *Osm*-sufficient animals with colitis and are low in *Osm*-deficient mice (*36*).

In our system, we did not observe lower viral titers in *Osm*-deficient mice, despite increased levels of IFN-I. This is likely due to the genetic background of mice used in our study. Inbred mouse strains, including C57BL/6, lack functional alleles of the *Mx1* gene. Mx1 is the key ISG with anti-influenza activity via blockade of viral transcription (*39, 40*), therefore IFN-I plays a minor role in antiviral defense in *Mx1*-deficient mice (*41*). Thus, in the absence of Mx1, the balance of IFN-I responses during influenza is shifted from defensive towards pathological.

Together with recent studies (*9, 24, 29*), our findings emphasize the pathological potential of unrestrained IFN-I responses. Macrophage-derived OSM exemplifies a host mechanism to control IFN-I in the context of viral stimuli to further protect the infected host. Furthermore, our work elucidates an anti-inflammatory macrophage-epithelial cell communication circuit that may be generalizable in other tissue sites where OSM is induced during inflammation. As evident from the recent SARS-CoV-2 pandemic, emerging and re-emerging respiratory viruses pose a significant ongoing threat to human health. In fact, severe disease in COVID-19 patients often results from uncontrolled immune responses as opposed to uncontrolled viral replication (*42*–*46*). Therefore, understanding the intricate regulatory mechanisms of IFN-I may represent a novel pan-viral strategy to dampen immune responses and promote the restoration of tissue homeostasis following respiratory infection.

## Acknowledgments

We thank all members of the Franklin and Medzhitov laboratories for thoughtful discussions and C. Zhang, C. Annicelli, and S. Cronin for supporting the Medzhitov lab animal colony. We thank C. Benoist for guidance regarding RNA-sequencing and A. Wang, R. Jackson, and I. Zanoni for critical feedback on the manuscript.

## Funding

National Institutes of Health grant 1R35GM150816-01 (RAF)

National Institutes of Health grant 5P30DK043351-32 (RAF)

National Institutes of Health grant 5TL1TR002543-04 (PRM)

National Science Foundation Graduate Research Fellowship Program grant DGE1122492 (PRM)

Cancer Research Institute Irvington Donald J. Gogel postdoctoral fellowship (RAF)

Howard Hughes Medical Institute (RM)

## Author contributions

Conceptualization: RAF, RM

Formal analysis: DAH, PR-M, AOM, RAF, SDP, JL Funding acquisition: RAF, RM

Investigation: DAH, PR-M, AOM, RAF, SY, AL, ABV

Resources: SL, XZ, MOL (*Fcgr1*^Cre^ mice)

Visualization: DAH, PR-M, AOM, RAF

Writing – original draft: DAH, PR-M, AOM, RAF

Writing – reviewing and editing: DAH, PR-M, AOM, RAF, RM

## Data and materials availability

Sequencing data are available from the National Center for Biotechnology Information Gene Expression Omnibus (GSE #249668). All data are available in the main text or the supplementary materials.

## Supplementary Materials

Materials and Methods

Figures S1 to S10

Tables S1 to S2

