## Supplemental materials for "Macrophages control pathological interferon responses during viral respiratory infection"

**The PDF file includes:**

Materials and Methods
Figures S1 to S10
Tables S1 to S2

**Materials and Methods**

Mice

All animal experiments were performed in accordance with institutional regulations after protocol review and approval by the Institutional Animal Care and Use Committee (IACUC) at Harvard Medical School. C57BL/6J mice (stock no. 000664) were obtained from Jackson Laboratories (JAX), *Fcgr1*$^{Cre}$ mice were provided by Dr. Ming Li (MSKCC), and gene targeted Osm "Knockout-first" mice were obtained from the KOMP repository (ES cell line Osmtm1a(KOMP)Wtsi). To generate conditional *Osm* alleles, mice were bred to the FLPo deleter strain (stock no. 012930; JAX). *Osm* conditional mice were then bred to *Fcgr1*$^{Cre}$ to achieve conditional deletion. Female mice aged 8 to 12 weeks were used for all experiments unless otherwise noted. Mice were housed under specific pathogen-free conditions.

Infection and *in vivo* treatments

For influenza virus infection, mice were anesthetized with a ketamine/xylazine mixture and 225 - 300 plaque forming units (pfu) of virus were administered intranasally in a 30µl volume. For intratracheal treatments, mice were anesthetized by inhalation of isoflurane/propanediol. 50µg of HMW poly(I:C) (InvivoGen) in a 50µl volume of normal saline was administered using a pipette. For IFNAR-1-blockade treatments, mice were injected intravenously by retro-orbital injection with 200µg of *InVivo*Plus anti-mouse IFNAR-1 or rat IgG2a isotype control (Bio X Cell) in 100µl volume. For murine rOSM rescue experiments, mice were administered 1µg of murine rOSM (Bio-Techne) in 10µL or PBS in addition to daily treatment with 50µg of HMW poly(I:C). Body weights were monitored daily. For the Evans blue dye (EBD) lung barrier permeability assay, mice were injected intraperitoneally with 10mg/kg of EBD (Sigma) 16 hours before euthanasia.

Plaque assay and virus propagation

Influenza virus strain A/WSN/33 was originally obtained from Dr. Peter Cresswell (Yale University School of Medicine). The virus was propagated and titered by plaque assay using Madin-Darby Canine Kidney (MDCK) cells (ATCC). To determine viral titer, a 10-fold serial dilution of virus was prepared in in 0.1% BSA in PBS in 200µl volume. The cells were infected at 37°C for 1 hour and shaken every 15 minutes. Cells were washed twice with PBS and an agarose overlay of 1% agarose gel mixture including TPCK Trypsin (1:2000), 0.21% BSA, 0.225% $NaHCO_3$, 100U/mL penicillin/streptomycin in MEM was added to each well and the plate was incubated at 37°C for 48 hours upside down. On day 3 the gel was removed and 1mL of crystal violet solution (Sigma-Aldrich) was added to each well, and the plate was incubated at room temperature for 30 minutes. For viral propagation, MDCK cells were infected at an MOI of 0.002 in 100µl of DMEM + 1% BSA per well and incubated for 10 minutes at 37°C. 1.5mLs of DMEM +1% BSA were added to each well and cells were left to grow for 48 hours. Cells were spun down at 2000 RPM for 10 minutes and supernatant harvested.

Animal harvest and cell isolation

Animals were euthanized by administration of a lethal dose of ketamine/xylazine mixture and perfused with 2mM EDTA in PBS. The trachea was exposed, nicked, and a 22g catheter was inserted. A 1mL-PBS-filled syringe was attached to the catheter and used to inflate then deflate lungs to collect cells present in BALF. The syringe was replaced with a 1mL-dispase-filled syringe and lungs were inflated. Inflated lungs were removed, and lung lobes were chopped and immersed in enzyme solution (100µg/mL DNAse and 83µg/mL Liberase in RPMI) and placed in a 37°C shaker for 40 minutes. BALF and lung digest cells were filtered through a 70µm cell strainer to obtain a single cell suspension and exposed to hypotonic lysis (ACK lysing buffer, ThermoFisher) to remove red blood cells.

Flow cytometry and cell sorting

Antibodies used for flow cytometry are listed in Table S1. Cell viability was determined using Zombie Aqua Fixable Viability Kit (BioLegend) following manufacturer's protocol for flow cytometry and using DAPI at 10.9µM (BioLegend) for cell sorting. Samples were Fc-blocked with purified anti-mouse CD16/32 antibody (BioLegend). For surface staining analysis, cells were fixed in 2% PFA for 20 minutes at room temperature then washed. For pSTAT3 staining, following fixation, samples were washed and permeabilized overnight in 90% methanol at -20°C. After permeabilization, samples were washed with ice-cold FACS buffer and stained for pSTAT3 and any surface antigens that were marked with methanol-sensitive fluorophores. Flow cytometry samples were acquired on a Symphony A5 or A1 (BD) and analyzed using FlowJo (Tree Star Technologies). Sorting samples were acquired in an Aria 561 (BD) or MoFLo Astrios (Beckman Coulter Life Sciences).

Blood plasma and BALF collection and protein analysis

Blood was harvested through retro-orbital bleeding and plasma was isolated using lithium heparin coated plasma separator tubes (BD). Total BALF protein levels were quantified using the Bio-Rad Protein Assay. Mouse IFN-α2 was quantified using the LumiKine Xpress mIFN-α 2.0 kit (Invivogen) and mouse IFN-β quantified using the LumiKine Xpress mIFN-β 2.0 kit (Invivogen) according to manufacturer instructions. Mouse OSM was quantified using the Mouse OSM Quantikine ELISA kit (R&D Systems) according to manufacturer instructions. For EBD barrier permeability assay, 200µl of collected BALF was read at 620nm absorbance in a 96-well plate. All plates were read using a BioTek Synergy HTX plate reader.

RNA Extraction and Quantification

Tissue samples were collected into TRIzol reagent (Invitrogen) and homogenized in microtubes with ceramic beads using a Bead Mill Homogenizer (OMNI International) and RNA was extracted using the Direct-zol RNA Miniprep kit (Zymo Research). RNA from *in vitro* samples was extracted using phenol-chloroform isolation. For sorted cells, samples were collected into TRIzol LS reagent (Invitrogen) and RNA was harvested via phenol-chloroform isolation. RNA samples used for *Ifnb* or *Ifna4* RT-qPCR analysis were treated with DNA-free DNA Removal Kit (Invitrogen) before cDNA synthesis. cDNA synthesis was performed using MMLV reverse transcriptase (Takara Bio) and oligo(dT) primers. RT-qPCR reactions were performed on the Applied Biosystems QuantStudio5 Real-Time PCR System (ThermoFisher) using PowerUp SYBR Green Master Mix (ThermoFisher). Relative expression in RT-qPCR analysis was calculated as $(2^{-\Delta Ct})$*1000. Fold induction in RT-qPCR analysis was calculated as $(2^{\wedge -ddCt})$, where dCt values were normalized to wild-type controls. ΔCt was calculated by subtracting *Rpl13* Ct values for murine cells or *TUB1A1* Ct values for human cell lines from Ct values of genes of interest. All oligonucleotides are listed in Table S2.

Cell culture

To generate bone-marrow derived macrophages (BMDMs), animals were euthanized by administration of a lethal dose of ketamine/xylazine mixture, and their femurs and tibias were isolated and cleansed with ethanol. Bone marrow was flushed with RPMI 1640 and cells were treated with ACK lysis buffer. Cells were cultured in complete RPMI (RPMI 1640 +2mM L-glutamine, 1mM sodium pyruvate, 10mM HEPES, 200U/mL penicillin/streptomycin, 10% FBS, and 2-mercaptoethanol (BME)). The following day (day 1), non-adherent cells were harvested and $10x10^6$ cells were resuspended in a mixture of 70% complete RPMI and 30% L929-conditioned media and plated on a petri-dish. On day 4, growth media was supplemented. Cells were used for experiments on day 6. All cell cultures were maintained in a 37°C incubator at 5% $CO_2$. Stress conditions were as follows: ER stress was induced by 5µm thapsigargin, heat shock performed at 42°C, 10ng/mL LPS, and 50µg/mL poly(I:C). MDCK cells (ATCC) were cultured in complete DMEM (2mM L-glutamine, 1mM sodium pyruvate, 10mM HEPES, 200U/mL penicillin/streptomycin, and 10% FBS). Adenocarcinoma human alveolar basal epithelial cells (A549) were obtained from ATCC (CCL-185) and cultured in DMEM supplemented with 200U/mL penicillin/streptomycin and 10% FBS. A549s were treated with 20ng/mL of human rOSM (Bio-Techne) 12 hours before transfection with 50µg of poly(I:C) HMW (Invivogen) using Lipofectamine RNAiMax (Invitrogen).

Bulk RNA sequencing

For tissue samples, RNA was extracted as previously described. Purified RNA samples were diluted to 2ng/µl, and 1µl of the sample was suspended in 5µL of TCL buffer with 1% BME (Sigma-Aldrich). Data was processed for sequencing and normalization according to the Immgen protocol (https://www.immgen.org/img/Protocols/ImmGenULI_RNAseq_methods.pdf).

Transcriptional Analysis

Normalized reads were quality filtered based on minimum expression and coefficient of variation, and then visualized with volcano plots and FC/FC plots using Multiplot Studio (GenePattern; Broad Institute). Pathway analysis was performed using Enrichr (*20*). Heat map visualizations were performed using Morpheus (Broad Institute, https://software.broadinstitute.org/morpheus).

Statistical Analysis

Results were statistically analyzed using a Student's t test or two-way ANOVA as appropriate using Prism 10.0 (GraphPad Software, Inc). If a difference in variances between groups was determined, a Mann-Whitney U test was used. Kaplan-Meier survival curves were compared using a log-rank Mantel-Cox test.

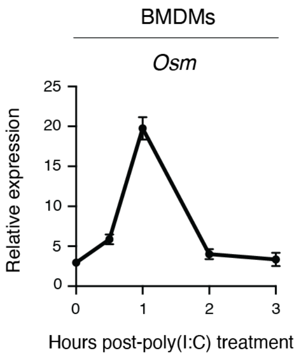


**Fig. S1. OSM is induced by viral stimuli.** BMDMs were treated with poly(I:C) at 50µg/mL and RNA collected at indicated time points (n = 3). Representative of two independent experiments. Relative expression calculated as $(2^{-\Delta Ct})*1000$.

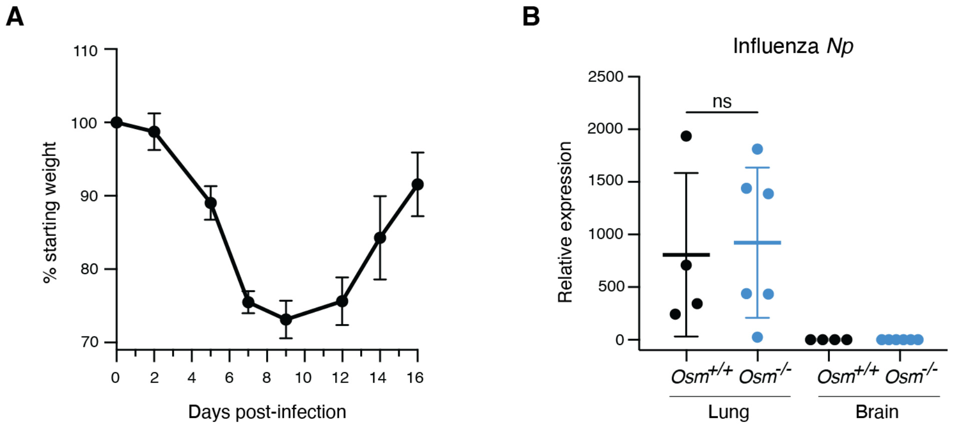


**Fig. S2. Wild-type mice recover from sublethal IAV infection and IAV RNA is not disseminated in *Osm*$^{-/-}$ mice.** (**A**) Mice were infected intranasally with 300 PFU of A/WSN/1933(H1N1) and monitored to assess weight loss (n = 4). (**B**) RT-qPCR analysis of IAV transcripts encoding nucleoprotein (*Np*) in whole lung or brain homogenate at 6dpi (n = 4 - 6). All data are representative of at least two independent experiments. Relative expression calculated as ($2^{-\Delta Ct}$)*1000. Error bars indicate SD; ns = not significant.

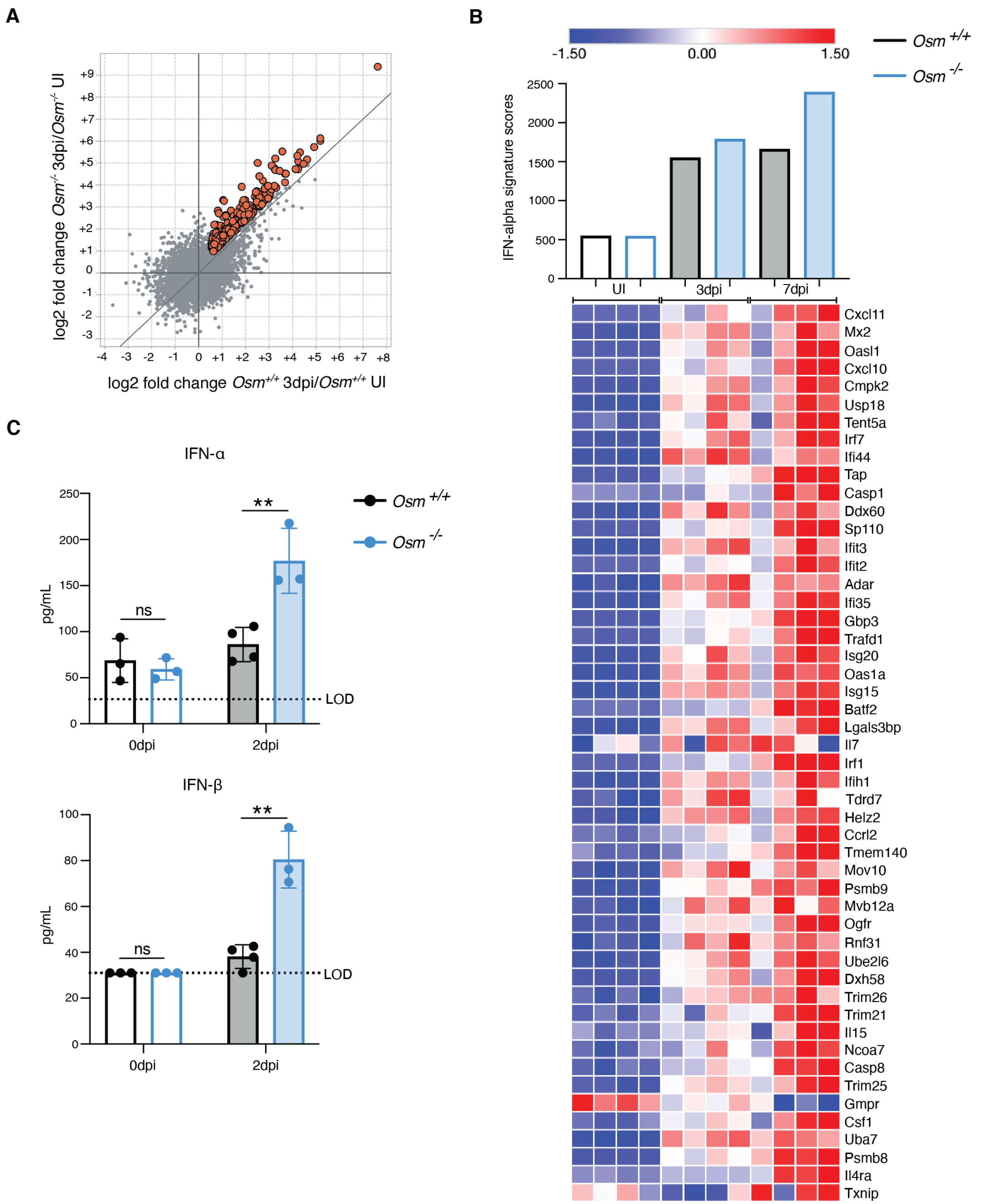


**Fig. S3. *Osm*$^{-/-}$ mice exhibit an enhanced IFNα response during IAV infection.** (**A**) FC/FC plots comparing gene expression values at 3dpi versus UI, in *Osm*$^{-/-}$ mice versus *Osm*$^{+/+}$ (n = 2). Highlighted genes were induced at least 1.5x fold-change in 3dpi/UI lungs for both *Osm*$^{+/+}$ and

*Osm*$^{-/-}$ mice and had 1.25x higher fold change values in *Osm*$^{-/-}$ compared to *Osm*$^{+/+}$ mice. (**B**) Expression of MSigDB hallmark IFNα response signature genes in whole lung. Average expression of IFNα response genes (top) and heatmap of selected transcripts from the MSigDB hallmark IFNα response signature gene set (bottom). The top 50 genes that displayed enhanced upregulation in *Osm*$^{-/-}$ mice at 3dpi, as identified in fig. S3A, were selected for display. Color scale reflects row Z-score values. UI = uninfected. (**C**) Mice were infected intranasally with 225 PFU of A/WSN/1933(H1N1), and protein from BALF was harvested at indicated time points (n = 3 - 4; representative of two independent experiments). IFN-α ELISA for IFN-α2 subtype. Error bars indicate SD, $^{**}p \leq 0.01$ ns = not significant, (unpaired Student's t test for (C)).

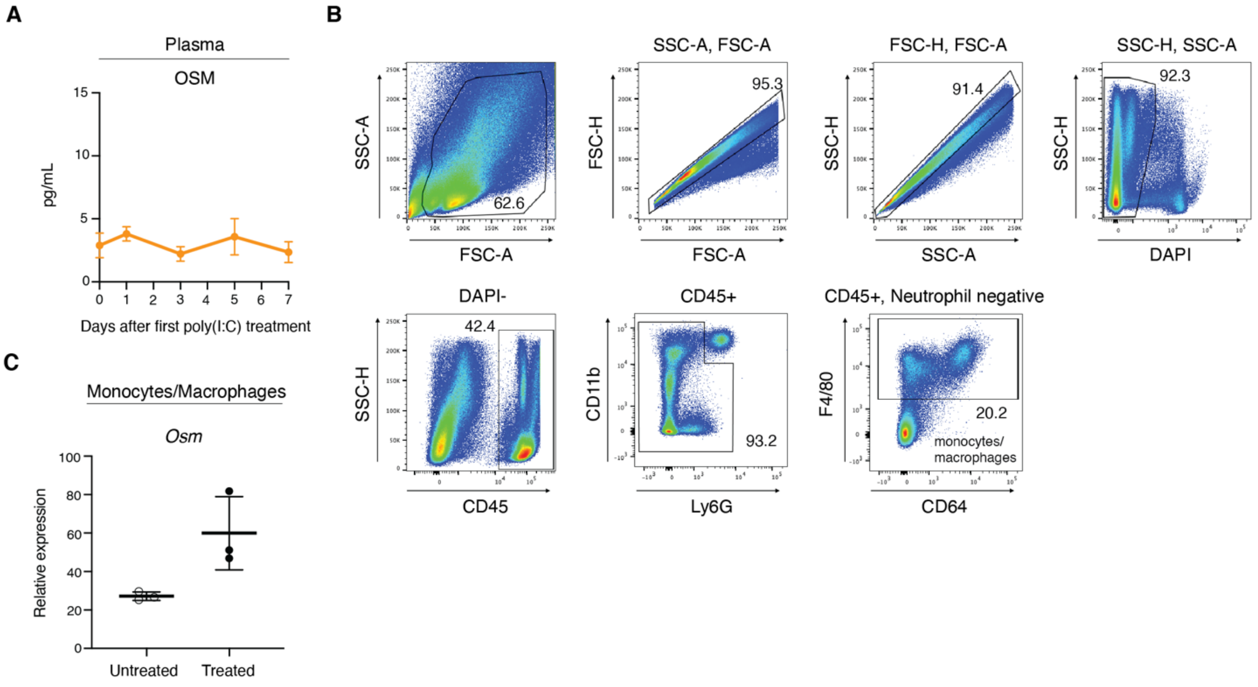


**Fig. S4. Following poly(I:C) treatment OSM is not detectable in plasma but is induced in lung monocytes and macrophages.** (**A**) Mice were treated i.t. with poly(I:C) (50µg) daily for five consecutive days. Plasma was collected for analysis of OSM protein levels at the indicated time-points (n = 4 - 5; representative of two independent experiments). (**B**) Representative gating strategy for FACS of monocytes/macrophages from digested lungs ($CD45^+$ $F4/80^+$ viable singlets). (**C**) RT-qPCR quantification of *Osm* expression in sorted monocytes/macrophages (n = 3). Relative expression calculated as $(2^{-\Delta Ct})*1000$.

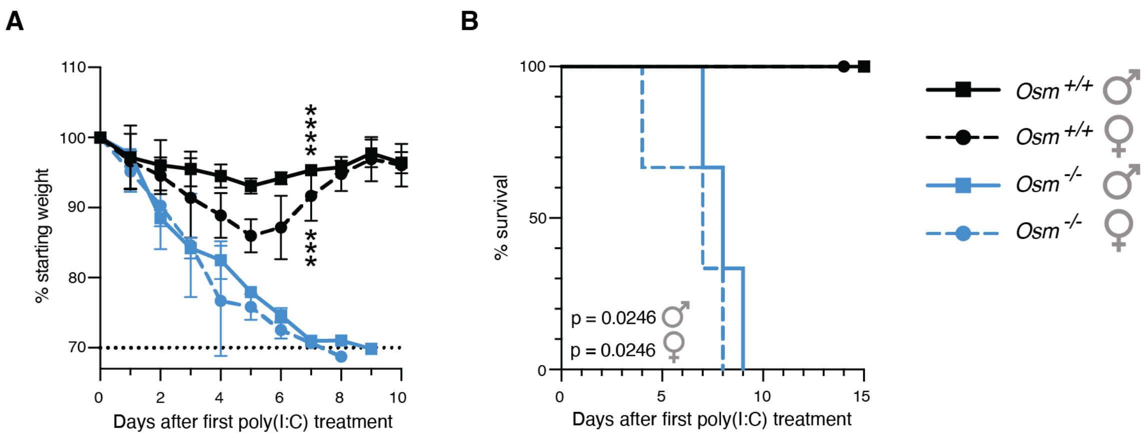


**Fig. S5. Susceptibility to poly(I:C)-induced mortality in *Osm*$^{-/-}$ mice is conserved across sexes. (A and B)** Male and female mice were treated i.t. with poly(I:C) (50µg) daily for five consecutive days and monitored daily for percent initial body weight (A) and survival (B) (n = 3; representative of two independent experiments). Error bars indicate SD, ***$p \leq 0.001$, ****$p \leq 0.0001$ (two-way ANOVA up to day 7 for (A), log-rank Mantel-Cox test for (B).

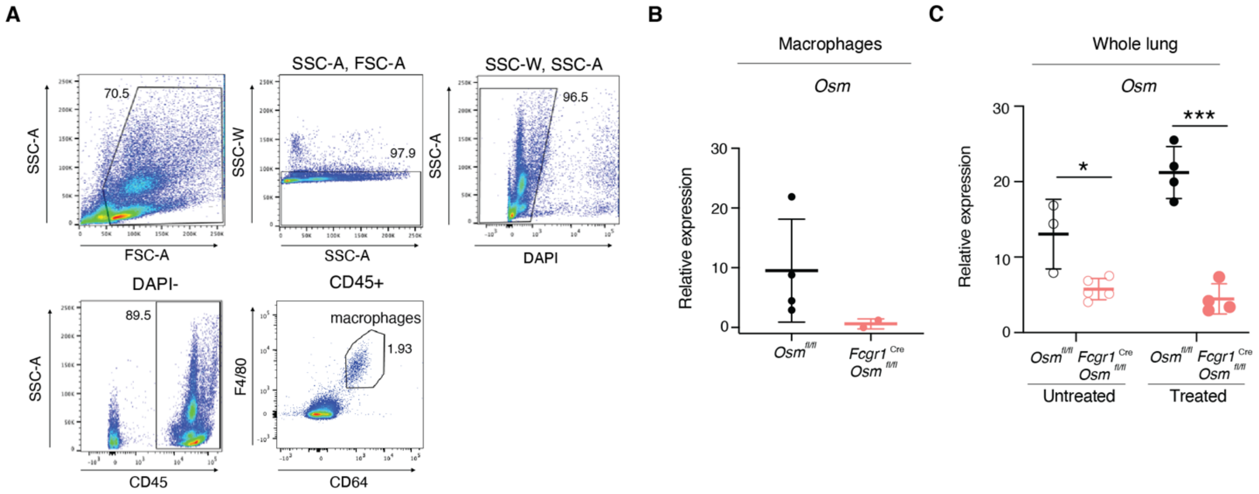


**Fig. S6. Analysis of *Osm* expression in macrophages from *Fcgr1*$^{\mathrm{Cre}}$*Osm*$^{\mathrm{fl/fl}}$ mice.** (**A**) Representative gating strategy for FACS of macrophages from digested lungs in mice at baseline ($CD45^{+}$ $CD64^{+}$ $F4/80^{+}$ viable singlets). (**B**) RT-qPCR quantification of *Osm* transcripts in sorted macrophages from *Fcgr1*$^{\mathrm{Cre}}$*Osm*$^{fl/fl}$ and *Osm*$^{fl/fl}$ mice (n = 3). (**C**) *Fcgr1*$^{\mathrm{Cre}}$*Osm*$^{fl/fl}$ and *Osm*$^{fl/fl}$ mice were treated i.t. with poly(I:C) (50µg) daily for five consecutive days. Whole lung was collected at baseline and at day 7 after the first poly(I:C) treatment. RT-qPCR quantification of *Osm* transcripts in whole lung (n = 3 - 5). Relative expression calculated as $(2^{-\Delta Ct})*1000$. Error bars indicate SD, *$p \leq 0.05$, ***$p \leq 0.001$, (unpaired Student's t test or Mann-Whitney U test for (C)).

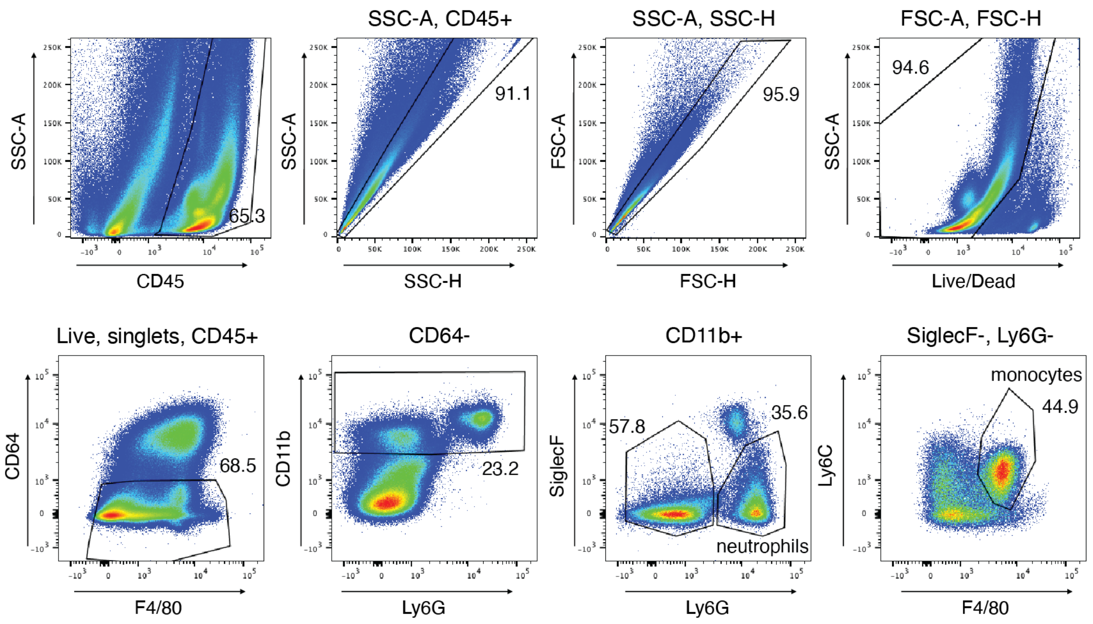


**Fig. S7. Analysis of monocyte and neutrophil infiltration by flow cytometry.** Representative gating for analysis of monocyte ($CD45^+$ $CD64^-$ $CD11b^+$ $SiglecF^-$ $Ly6G^-$ $Ly6C^+$ $F4/80^+$ viable singlets) and neutrophil ($CD45^+$ $CD64^-$ $CD11b^+$ $SiglecF^-$ $Ly6G^+$ viable singlets) lung infiltrates in *Fcgr1*$^{Cre}$*Osm*$^{fl/fl}$ and *Osm*$^{fl/fl}$ poly(I:C)-treated mice.

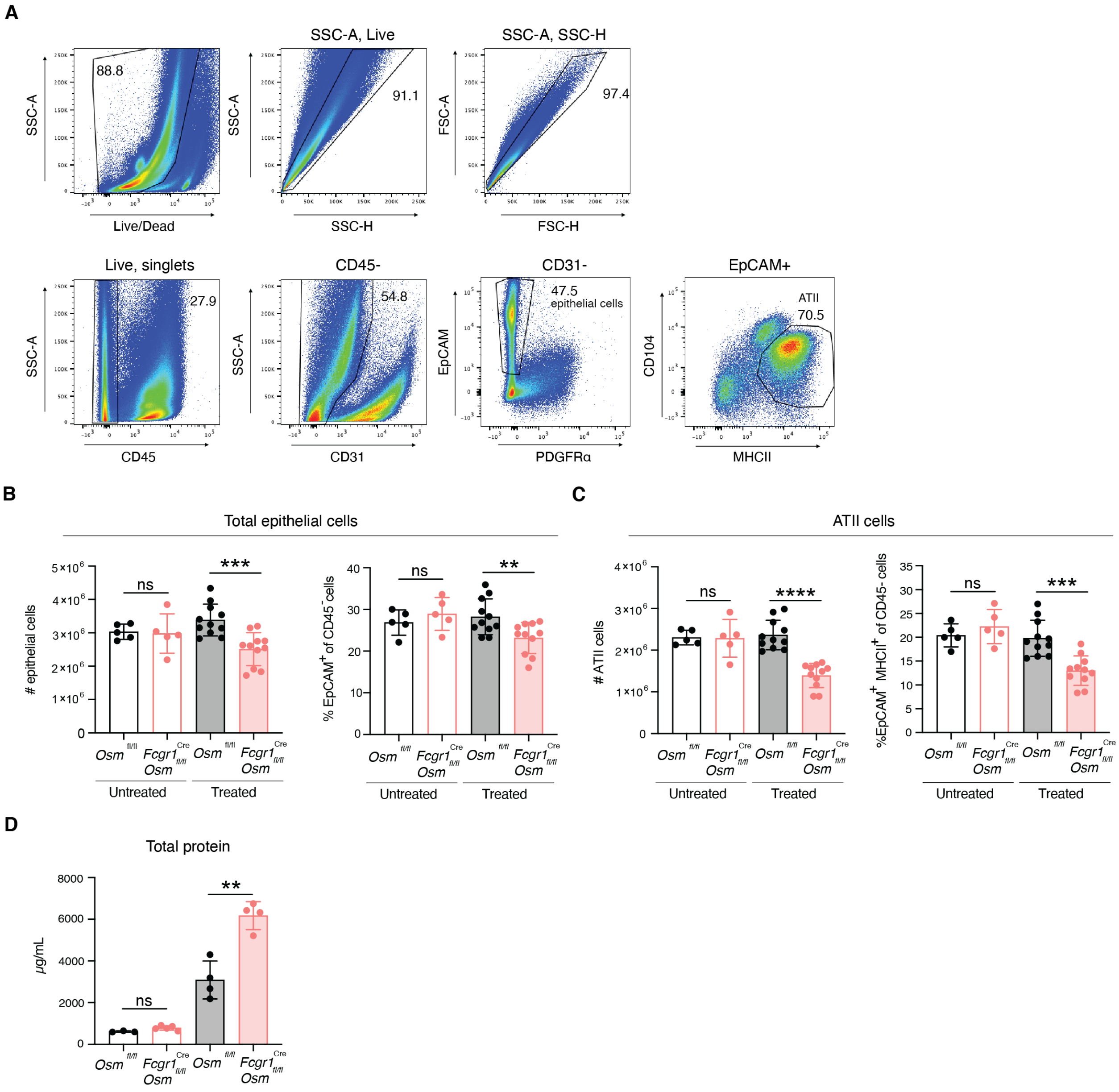


**Fig. S8. Analysis of epithelial barrier damage in poly(I:C) injured lungs.** (**A**) Representative gating for total epithelial cells (CD45$^-$CD31$^-$ PDGFRα$^-$ EpCAM$^+$ viable singlets) and type II alveolar epithelial cells (ATIIs) (CD45$^-$ CD31$^-$ PDGFRα$^-$ EpCAM$^+$ MHCII$^{high}$ viable singlets). (**B, C, and D**) *Fcgr1*$^{Cre}$*Osm*$^{fl/fl}$ and *Osm*$^{fl/fl}$ mice were treated i.t. with poly(I:C) (50µg) daily for five consecutive days. Lungs were harvested and digested for analysis of total epithelial cell loss (B), ATII loss (C), and total protein accumulation in BALF at baseline and at day 7 after the first poly(I:C) treatment (D) (n = 5 - 11; pooled from three independent experiments for poly(I:C) treated mice; total protein quantification in BALF (n = 3 - 4). Error bars indicate SD, $^{**}p \leq 0.01$, $^{***}p \leq 0.001$, $^{****}p \leq 0.0001$, ns = not significant, (unpaired Student's t test for (B), (C), (D)).

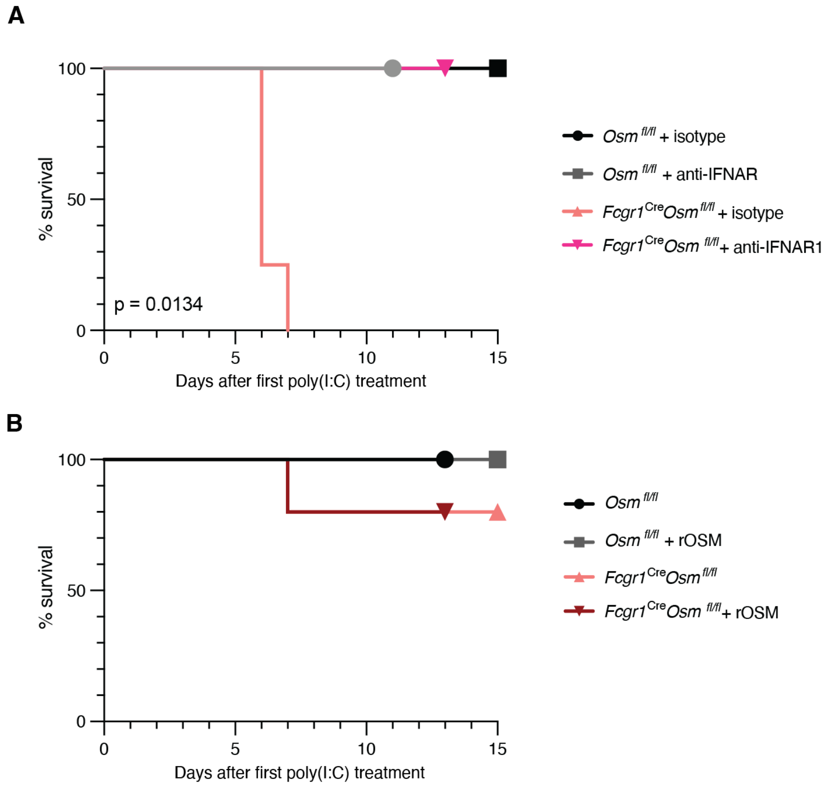


**Fig. S9. Survival curves of mice rescued with anti-IFNAR1 blockade or local rOSM treatment.** (**A**) Mice were treated i.t. with poly(I:C) (50µg) daily for five consecutive days. In addition, mice were treated i.v. with either 200µg of anti-mouse IFNAR-1 or isotype control antibody diluted in 100µl PBS on day -1, day 0, day 1, day 3, and day 4 after the first poly(I:C) treatment. Mice were monitored daily (n = 3 - 4). (**B**) Mice were treated i.t. with 1µg of murine rOSM in addition to 50µg of poly(I:C) daily for five consecutive days and monitored daily (n = 4 - 5). All data are representative of at least two independent experiments. Log-rank Mantel-Cox test.

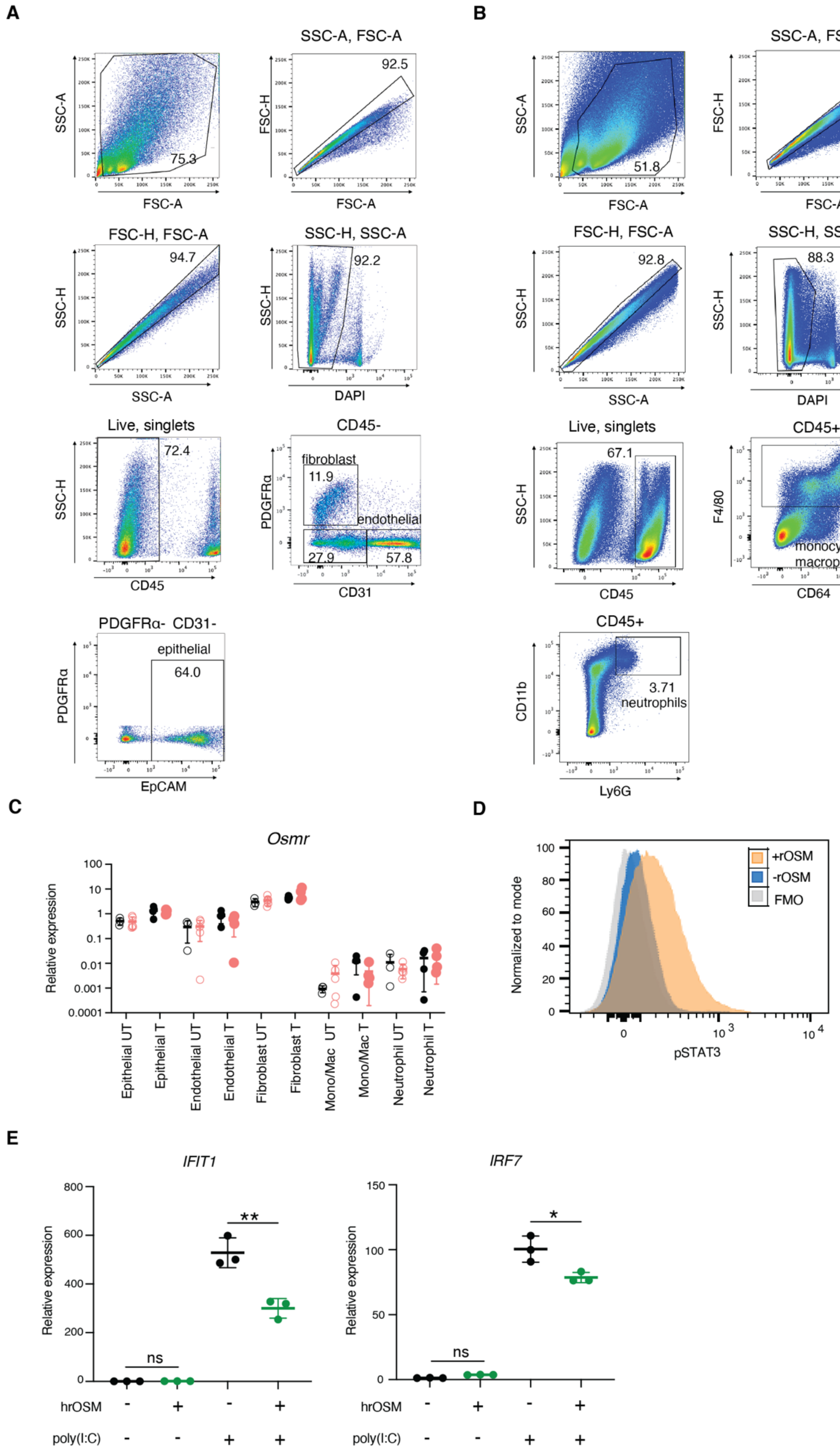

A
SSC-A
FSC-A
75.3
SSC-A, FSC-A
FSC-H
92.5
FSC-H, FSC-A
SSC-H
94.7
SSC-H, SSC-A
92.2
DAPI
Live, singlets
72.4
CD45
CD45-
PDGFRα
fibroblast
11.9
endothelial
27.9
57.8
CD31
PDGFRα- CD31-
epithelial
64.0
EpCAM
B
51.8
94.1
92.8
88.3
67.1
CD45+
F4/80
24.8
monocytes/
macrophages
CD64
CD11b
3.71
neutrophils
Ly6G
C
Osmr
Relative expression
Epithelial UT
Epithelial T
Endothelial UT
Endothelial T
Fibroblast UT
Fibroblast T
Mono/Mac UT
Mono/Mac T
Neutrophil UT
Neutrophil T
D
Normalized to mode
+rOSM
-rOSM
FMO
pSTAT3
E
IFIT1
IRF7
ns
**
*
hrOSM
poly(I:C)

**Fig. S10. Epithelial cells respond to OSM.** (**A**) Representative gating to obtain epithelial cells, endothelial cells, and fibroblasts is shown. Epithelial cells were identified as $CD45^-$ $PDGFR\alpha^-$ $CD31^-$ $EpCAM^+$, endothelial cells as $CD45^-$ $PDGFR\alpha^-$ $CD31^+$, and fibroblasts as $CD45^-$ $PDGFR\alpha^+$, all gated on viable singlets. (**B**) Representative gating to obtain neutrophils and macrophages/monocytes is shown. Neutrophils were identified as $CD45^+$ $CD11b^+$ $Ly6G^+$ and macrophages/monocytes were identified as $CD45^+$ $F4/80^+$, all gated on viable singlets. (**C**) Epithelial cells, endothelial cells, fibroblasts, macrophages, and neutrophils were sorted and RT-qPCR was performed for *Osmr* (n = 3 -5). (**D**) Cells isolated from digested lungs from WT mice were *ex vivo* stimulated with rOSM (100ng/mL) for 30 minutes and stained for surface markers and intracellular pSTAT3. Representative plot for pSTAT3 staining in epithelial cells following stimulation. Epithelial cells were defined as $EpCAM^+$ $CD45^-$ $CD31^-$ (n = 5; representative of two independent experiments). (**E**) A549 cells were treated with 20ng/mL of human rOSM for one hour and then transfected with 0.5µg of poly(I:C). Cells were harvested for RNA 18 hours after transfection. RT-qPCR was performed for *IFIT1* and *IRF7* (n = 3). Error bars indicate SD, $*p \leq 0.05$, $**p \leq 0.01$, ns = not significant, (unpaired Student's t test for (E)). Mono/Mac, monocytes and macrophages.

**Supplementary Tables**

Table S1: Flow cytometry antibodies

| Cell surface marker | Clone | Fluorochrome | Concentration | Provider |
|---|---|---|---|---|
| CD45 | 30-F11 | APC/Cy7 | 1:200 | Biolegend |
| CD45 | 30-F11 | APC | 1:200 | Biolegend |
| CD45 | I3/2.3 | FITC | 1:200 | Biolegend |
| CD64 | X54-5/7.1 | APC | 1:200 | Biolegend |
| CD64 | X54-5/7.1 | PE | 1:200 | Biolegend |
| F4/80 | BM8 | BV605 | 1:100 | Biolegend |
| F4/80 | BM8 | APC | 1:100 | Biolegend |
| F4/80 | BM8 | BV711 | 1:100 | Biolegend |
| MerTK | DS5MMER | PE/Cy7 | 1:200 | eBioscience |
| Siglec F | 1RNM44N | PE | 1:200 | Biolegend |
| Ly6C | HK1.4 | BV605 | 1:500 | Biolegend |
| Ly6G | 1A8 | PerCP-Cy5.5 | 1:300 | Biolegend |
| CD11c | N418 | APC/Cy7 | 1:300 | Biolegend |
| CD11b | M1/70 | BV421 | 1:800 | Biolegend |
| CD11b | M1/70 | PE/Cy7 | 1:800 | Biolegend |
| I-A/I-E | M5/114.15.2 | BV421 | 1:400 | Biolegend |
| EPCAM/CD326 | G8.8 | PE | 1:200 | Biolegend |
| PECAM1/CD31 | 390 | FITC | 1:200 | Biolegend |
| CD104 | 346-11A | PerCP-Cy5.5 | 1:200 | Biolegend |
| PDGFR$\alpha$/CD140a | APA5 | PE/Cy7 | 1:200 | Biolegend |

Table S2: Oligonucleotides

| Primer | Sequence |
|---|---|
| mouse *Ifit1* forward | 5’ - CAACCATGGGAGAGAATGCTG - 3’ |
| mouse *Ifit1* reverse | 5’ - TGCATCCCCAATGGGTTCTT - 3’ |
| mouse *Irf7* forward | 5’ - CAGCGAGTGCTGTTTGGAGAC - 3’ |
| mouse *Irf7* reverse | 5’- AAGTTCGTACACCTTATGCGG - 3’ |
| mouse *Isg15* forward | 5’ - AAGCAGCCAGAAGCAGACTC - 3’ |
| mouse *Isg15* reverse | 5’ - TTAGGTCCCAGGCCATTGCT - 3’ |
| mouse *Ifnb1* forward | 5’ - GTCTCATTCCACCCAGTGCT - 3’ |
| mouse *Ifnb1* reverse | 5’ - CAGCTCCAAGAAAGGACGAA - 3’ |
| mouse *Ifna4* forward | 5’ - CCAGAGAGTGACCAGCATCTAC - 3’ |
| mouse *Ifna4* reverse | 5’ - AAGGCCCTCTTGTTCCCGAG - 3’ |
| human *IFIT1* forward | 5’ - TCGGAGAAAGGCATTAGATC - 3’ |
| human *IFIT1* reverse | 5’ - GACCTTGTCTCACAGAGTTC - 3’ |
| human *IRF7* forward | 5’ - GGTGTGTCTTCCCTGGATAG - 3’ |
| human *IRF7* reverse | 5’ - GCTCCAGCTCCATAAGGAAG - 3’ |
| human *TUBA1A* forward | 5’ - GCCTGGACCACAAGTTTGAC - 3’ |
| human *TUBA1A* reverse | 5’ - TGAAATTCTGGGAGCATGAC - 3’ |
| *NP* forward | 5′ - GACGATGCAACGGCTGGTCTG - 3′ |
| *NP* reverse | 5′ - ACCATTGTTCCAACTCCTTT - 3′ |
| mouse *Rpl13* forward | 5’ - AGTATCTGGCCTTTCTCCGG - 3’ |
| mouse *Rpl13* reverse | 5’ - CCGAACAACCTTGAGAGCAG - 3’ |
